# High resolution visual information via a gap junction-mediated spike order code

**DOI:** 10.1101/2020.08.14.250910

**Authors:** Rudi Tong, Stuart Trenholm

**Affiliations:** Montreal Neurological Institute, McGill University, Montreal, Canada

## Abstract

Gap junctions promote correlated spiking between coupled neurons. Recording from pairs of coupled retinal ganglion cells, we find that differences in firing rate between coupled neurons dictates which cell leads correlated spike-pairs, and that the precise temporal order of spike activity between pairs of cells changes depending on the spatial position of a visual stimulus. We thus demonstrate a spike order spatial code for encoding sensory information.

## Main Text

Throughout the CNS, neighbouring pairs of electrically coupled neurons exhibit fine-scale correlated spiking^1^, and the extent of correlated spiking is dynamically modulated by sensory inputs^2,3^. However, within a pair of coupled neurons, sensory stimulation can drive different levels of activity in each cell, but how such heterogeneous activity levels relate to correlated spiking remains unclear. In the retina, visual stimuli drive pairs of electrically coupled ganglion cells to fire correlated spikes within ~ 2 ms of one another^2^. However, due to the Gaussian nature and spatial offset of receptive fields of neighbouring ganglion cells^4^, most visual stimuli differentially activate neighbouring cells. The retina thus provides an ideal model for examining how stimuli that drive different levels of activity in neighbouring coupled cells modulate the dynamics of correlated spiking.

Upon recording light-evoked responses from pairs of electrically coupled retinal ganglion cells in response to a visual stimulus flashed over their shared receptive field position, and plotting cross-correlograms to examine correlated spiking, we noted a large variability in the appearance of the cross-correlograms (**Figure 1a**): some pairs exhibited symmetrical peaks of roughly equal amplitude on either side of the trough at 0 ms delay; other pairs exhibited asymmetric peaks in the cross-correlogram with the majority of correlated spikes being on a single side of 0 ms delay. This raises the possibility that for some pairs both cells are equally effective in driving a correlated spike in the neighbouring cell, whereas in other pairs one of the cells is more effective in driving synchronous spiking. However, such an interpretation is inconsistent with the symmetric junctional conductance and coupling coefficient that have been measured between coupled retinal ganglion cells^5,6^. Another possibility is that the visual stimulus was not perfectly centered over the shared receptive field location between the pair of cells, and thus in some cases more strongly drove one cell, with the difference in spike rates relating to the bias in which cell drove correlated spike-pairs.

**Figure 1.**
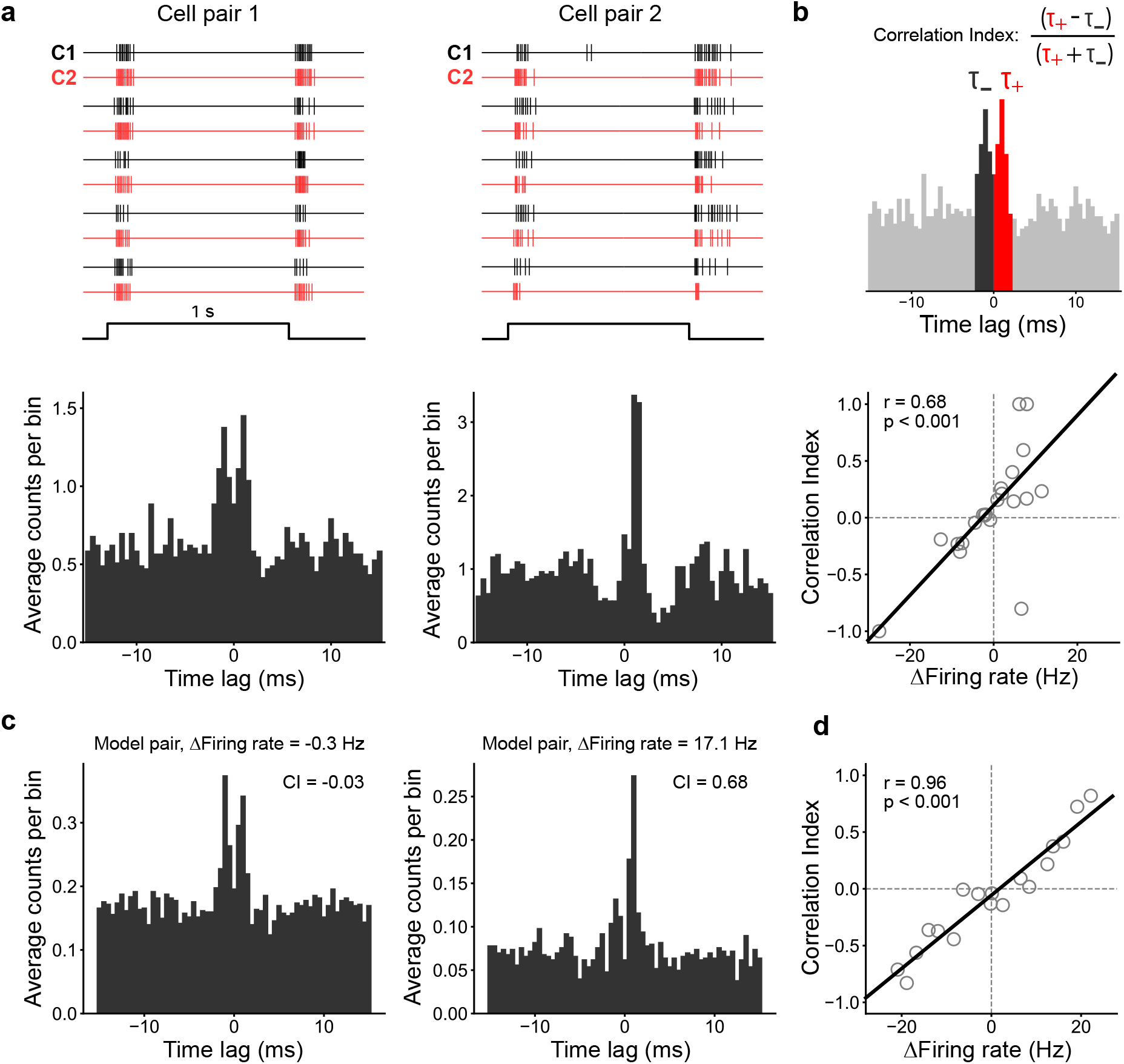
The difference in firing rate between coupled ganglion cells dictates which cell leads correlated spike-pairs. (**a**) *Top*, Example spike trains from two pairs of coupled ganglion cells (C1, cell 1, *black*; C2, cell 2, *red*), for five consecutive presentations of a visual stimulus flashed for 1 s over their shared receptive field location. *Bottom*, Cross-correlograms computed for the cell pairs outlined above. (**b**) *Top*, Explanation of the Correlation Index (CI; see Methods), defined as the normalized temporal delay between spike-pairs with positive (between 0 and 2 ms; red bins) and negative (between −2 and 0 ms; dark grey bins) lags. Bins beyond 2 ms delay are colored light grey. *Bottom*, A plot of the Correlation Index vs. absolute difference in spike rate between pairs of cells. The data (n = 21 cell pairs) was fit with a linear regression (r^2^ = 0.46). (**c**) Cross-correlograms of electrically coupled model neurons (see Methods) with equivalent (*left*) or different (*right*) firing rates. (**d**) Correlation Index calculated from a pair of model neurons as a function of firing rate difference between the cells (model parameters: spike transmission probability = 10 %, amplitude of depolarizing potential = ~6.5 mV; see Methods and **Supplementary Fig. 2**; r^2^ of linear regression = 0.92). In both (**b**) and (**d**), the listed r value is the Pearson correlation coefficient and the P-value is calculated with a permutation analysis of the Pearson correlation (see Methods).

We tested whether differences in light-evoked firing rates between pairs of coupled cells could account for biases in which cell drove correlated spike-pairs (and thus account for asymmetries in the cross-correlogram). To quantify biases in which cell drove correlated spike-pairs, we computed a Correlation Index (CI; **Figure 1b**, see Methods). For this metric, we first subtracted correlations expected purely by chance (via shuffling across trials and subtracting the ‘shift predictor’ calculated correlations; **Supplementary Figure 1**) and restricted our analysis to spike-pairs that occurred within a 2 ms window (though altering the size of the window used to define correlated spikes did not qualitatively affect the result; **Supplementary Figure 1**). CI values spanned from −1 to 1, with a CI value of −1 indicating that spikes in ‘Cell 1’ precede spikes in ‘Cell 2’ by 2 ms, and a CI value of 1 indicates that spikes in ‘Cell 2’ precede spikes in ‘Cell 1’ by 2 ms. We found a strong correlation between the difference in firing rate between a pair of cells and the measured CI value, such that the cell with the higher firing rate reliably led correlated spike-pairs (**Figure 1b**). It should be noted that while the ganglion cells we recorded from are ON-OFF cells^7^, and respond to both increments and decrements in light intensity, for all analyses we pooled ON and OFF responses together as both exhibited a similar relationship between Correlation Index and spike rate (**Supplementary Figure 1**).

To examine whether a difference in firing rate between a pair of coupled cells is sufficient to account for which cell led correlated spike-pairs, we generated a simplified computational model consisting of two pulse-coupled integrate-and-fire neurons (**Supplementary Figure 2**; see Methods). When the neurons were depolarized to mimic light-evoked spiking activity, the model cells exhibited correlated spiking, similar to what we found experimentally (**Figure 1c**; correlated spiking was absent when the model cells were uncoupled; **Supplementary Figure 2**). Introducing a difference in the spike rate between the two model cells was sufficient to bias which cell drove (i.e. led) correlated spike-pairs, with the cell with the higher firing rate leading correlated spike-pairs (**Figure 1d**; **Supplementary Figure 2**). As such, the difference in spike rate between pairs of coupled neurons appears to be sufficient to dictate which cell drives correlated spiking.

We hypothesized that since visual stimuli falling within different portions of the overlapping receptive field region between a pair of retinal ganglion cells will drive different levels of activity in both cells, the exact position of a stimulus should dictate which cell leads correlated spike-pairs. To test this, we flashed a small spot of light (40 μm diameter) at different locations over the shared receptive field region of pairs of coupled ganglion cells, and at each location we computed a cross-correlogram and the CI (**Figure 2a**). We found that progressively marching the spot of light across the shared receptive field region reliably modulated which cell led (i.e. drove) correlated spike-pairs (**Figure 2b**; each pair was stimulated at between 3-5 different spatial locations). The cell that drove the correlated spiking was consistently the cell with the higher firing rate (**Figure 2c**). Therefore, the timing and relative order of correlated spike pairs encodes high resolution spatial information.

**Figure 2.**
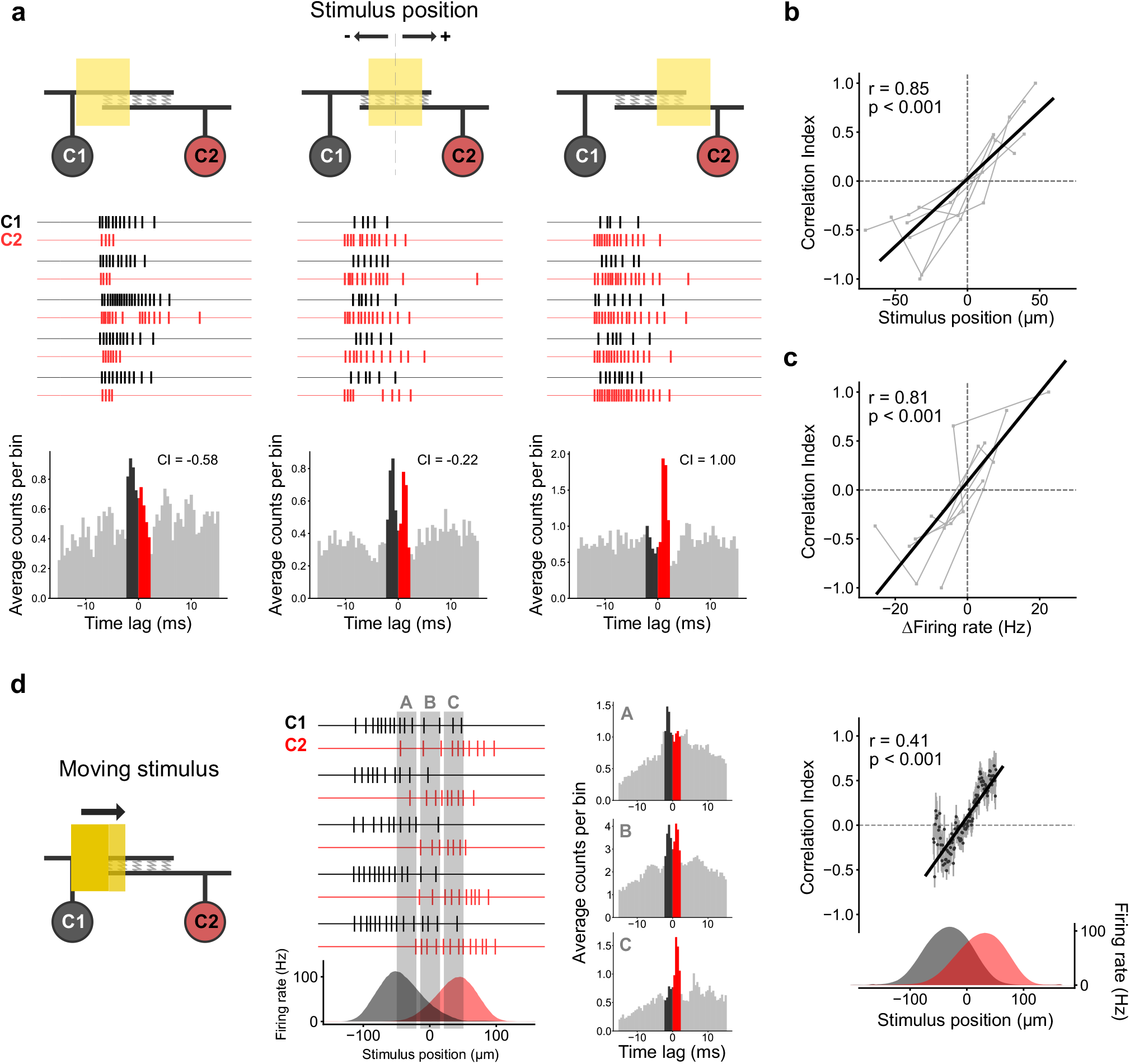
The relative order of correlated spike-pairs encodes high resolution spatial information. **(a)** Example spike trains are shown for a pair of cells presented with a small spot of light (40 μm diameter; 5 repetitions are shown; C1, *black*; C2, *red*) that was flashed at different locations within the region of receptive field overlap between a pair of coupled ganglion cells. Stimulus position is indicated in relation to the midpoint of the shared receptive field location between the cells (see schematic on top of the figure; see Methods). ON and OFF responses were pooled together (see Methods). A cross-correlogram is plotted for responses from each stimulus position, and the Correlation Index (CI) is indicated. **(b)** The Correlation Index is plotted as a function of stimulus position (n = 6 cell pairs; each cell pair was stimulated at between 3-5 different locations; r^2^ of linear regression = 0.72). **(c)** The Correlation Index is plotted as a function of firing rate difference between cells in each pair (n = 6 cell pairs; r^2^ of linear regression = 0.66). **(d)** *Left*, A 300 μm wide bar was moved over the receptive fields of a pair of coupled ganglion cells at 1000 μm/s along the preferred direction. *Middle*, Example spike trains (5 repetitions, top; C1, *black*; C2, *red*) and the average firing rate for a pair of coupled ganglion cells as the bar moved over their receptive fields (*bottom*; indicating the spatially-offset, Gaussian-like receptive fields of neighbouring cells; see Methods). Cross correlograms were computed for three different windows (A, B, C; indicated in grey) of the shared receptive field region of the pair of cells. *Right*, CI is plotted as a function of stimulus position (n = 11 cell pairs; **Supplementary Figure 3**). The data is fit with a linear regression (r^2^ = 0.16). Data points are shown as mean ± SEM. In (**b**-**d**), the listed r value is the Pearson correlation coefficient and the P-value is calculated with a permutation analysis of the Pearson correlation (see Methods).

As the coupled neurons we were recording from have previously been shown to be highly sensitive to visual motion (i.e. they are directionally selective ganglion cells^7,8^), we tested whether the spike order spatial code that we defined for static stimuli (**Figure 2b**) was also present for visual stimuli moving along the preferred direction. As a visual stimulus passed over the receptive fields of neighbouring cells, the relative order and timing of which cell led correlated spike-pairs progressed in a linear fashion (**Figure 2d; Supplementary Figure 3**), denoting that the spike order spatial code is accurately maintained during visual motion. Importantly, the spike order spatial code was invariant to changes in speed and contrast of a moving bar (**Supplementary Figure 3**), indicating the robustness of the code.

In summary, we show that fine-scale correlated spiking between pairs of coupled retinal ganglion cells is modulated by differences in their light-evoked firing rates, with the cell with the higher firing rate being more effective in driving (i.e. leading) synchronous spiking. We find that, due to the spatial offset of receptive fields of neighbouring coupled ganglion cells, combined with the Gaussian nature of their receptive fields, the relative timing and order of correlated spike-pairs accurately encodes the position of a visual stimulus.

The spike order code we present could be considered a version of rank order coding^9^, in which a decoder is tuned to prefer spikes from a group of input neurons arriving in a specific order. Modeling indicates that rank order coding could be a fast and efficient population coding strategy^9^, and such a coding scheme has been proposed in the retina^10^. In contrast to these population codes, our experiments reveal a spike order code that arises specifically between pairs of coupled neurons. While correlated spike-pairs from ganglion cells are thought to be particularly impactful in driving target neurons in retinorecipient regions^11^, it remains to be studied whether the particular spike order correlation code we present is processed by retinorecipient regions, though several previously described neuronal decoders could read out such a code^12–14^.

How does a difference in spike rate between coupled cells modulate which cell drives correlated spike-pairs? Previous work has shown that gap junctions enable correlated spiking between neighbouring ganglion cells via a complex interplay between chemical synaptic inputs, gap junction inputs, and dendritic non-linearities^2^: when one ganglion cell fires a spike, this generates a back-propagating action potential (bAP), which drives coupled spikelets in coupled post-junctional dendrites, and these spikelets combine with glutamatergic inputs from bipolar cells to drive dendritic spikes, which in turn drive correlated spikes^2^. In the present study, we elaborate upon this model: when there is a difference in firing rate between coupled cells, the cell with the higher firing rate generates more bAPs and thus drives more coupled spikelets in the post-junctional cell, therefore increasing the odds that the cell with the higher firing rate will drive correlated spiking (**Supplementary Figure 4**). As such, the described spike order code is critically dependent on the reciprocal nature of the gap junction connection. In the absence of gap junctional coupling, spike correlations between neighbouring ganglion cells are driven by common chemical synaptic input, which leads to correlated spiking on a timescale of ~100 ms and a symmetrical cross correlation centered around 0 ms delay^15^.

While, to our knowledge, this is the first description of a neuronal circuit that encodes information in the relative timing and order of correlated spike-pairs between coupled neurons, similar gap junction mediated fine-scale correlations – with peaks in the cross-correlogram around ± 2 ms, with a trough at 0 ms delay – have been described for pairs of coupled neurons in many other brain regions, including the inferior olive^16^, cerebellum^17^, and cortex^18^. As such, it is likely that a similar correlated spike order code is present in other brain regions, so long as during the course of being stimulated, pairs of electrically coupled cells are driven to different extents. The coupling-mediated spike order code we present might therefore represent a general coding strategy used throughout the CNS.

---

**Supplementary Figure 1.**
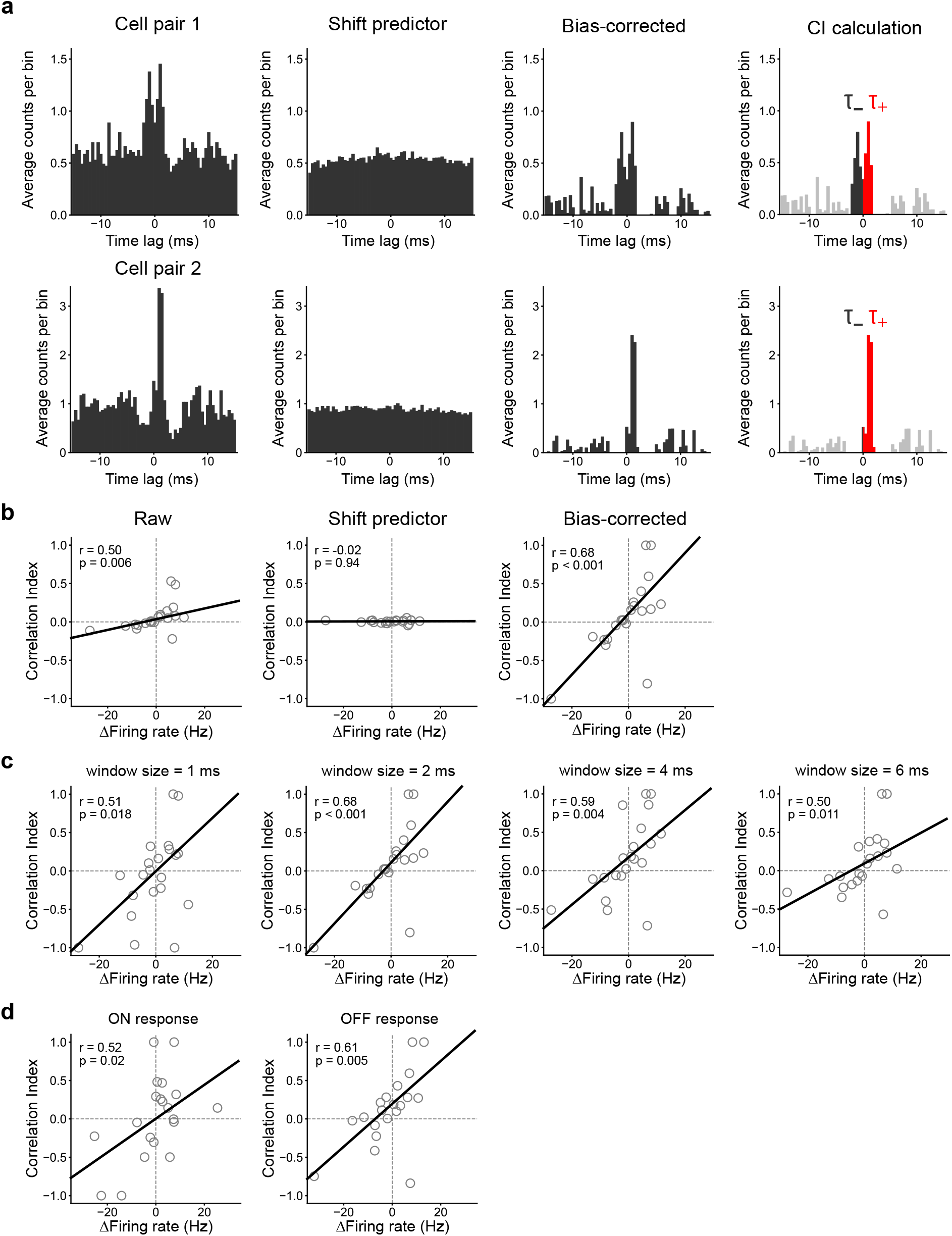
The Correlation Index as a read-out of the order and timing of correlated spike-pairs. **(a)** Examples of how the Correlation Index (CI) was calculated. From left to right, for two example cell pairs (*top*, *bottom*), the panels show the raw cross-correlogram, the shift predictor (computed across stimulus trials to correct for correlations expected by chance), the bias-corrected cross-correlogram (shift predictor subtracted from raw), and the spike-pairs used in the CI calculation (coloured in red and black). **(b)** *Left*, CI calculated for the ‘raw’ data before subtraction of the shift-predictor is plotted against ΔFiring rate between pairs of cells. *Middle*, CI is calculated for the shift-predictor data. *Right*, CI is calculated for the bias-corrected (shift-predictor subtracted) data. **(c)** Changing the window size which defines the maximum spike delay used in the calculation of the Correlation Index did not affect the finding that CI linearly varies with changes in the relative difference in spike rate between coupled cells. **(d)** Separated ON and OFF responses from the data that were pooled together in **Figure 1a,b**, showing the relationship between Correlation Index and ΔFiring rate is present for both ON and OFF responses.

**Supplementary Figure 2.**
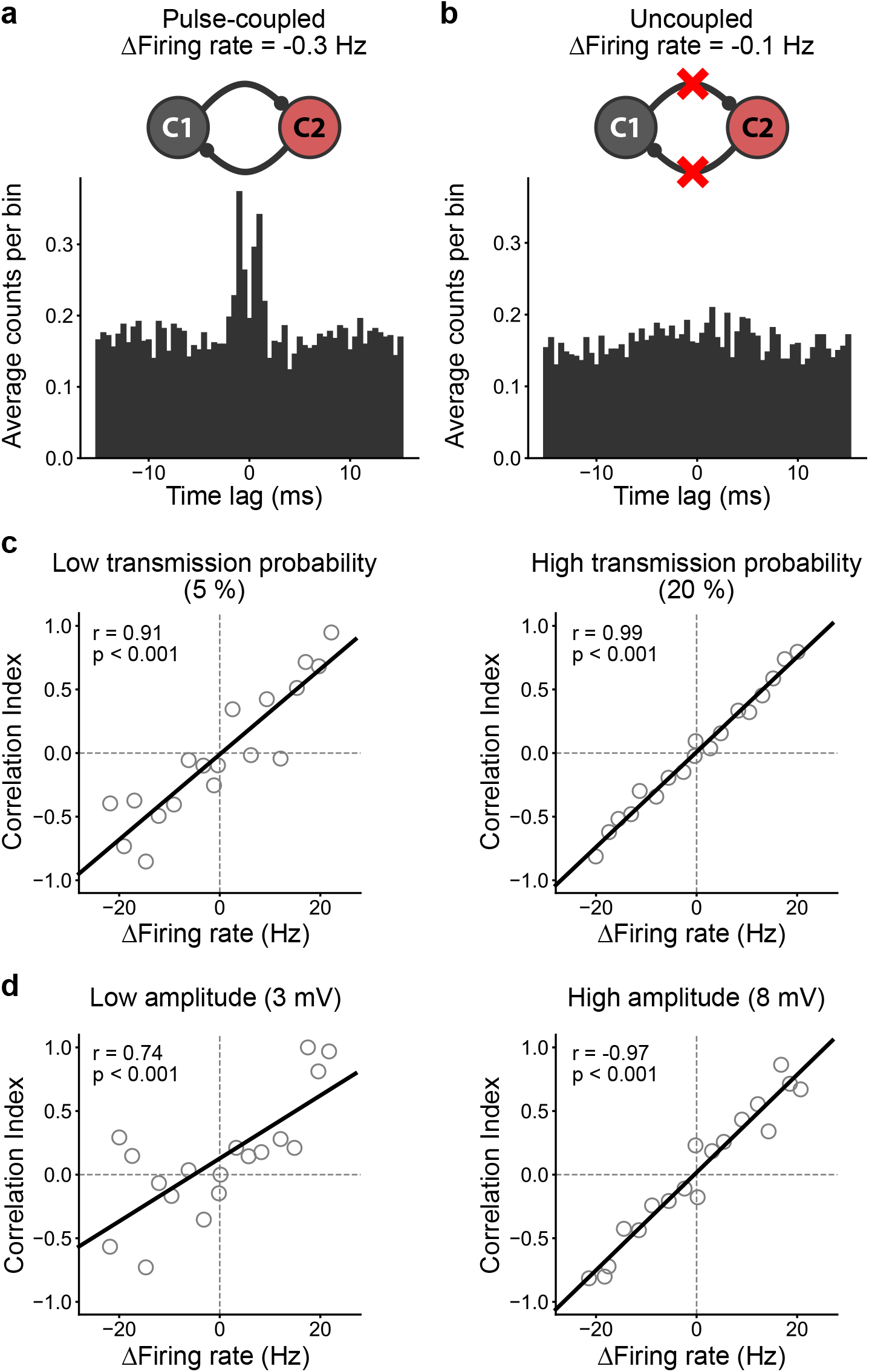
A pair of pulse-coupled integrate-and-fire neurons recapitulates the relationship between the Correlation Index and ΔFiring rate. **(a)** Example cross-correlogram of a pair of pulse-coupled model neurons (see Methods for details; C1, *black*; C2, *red*). The coupling was modelled as a pair of strong, probabilistic synapses, which, after the arrival of an action potential in one of the cells, leads to a depolarizing potential in the postjunctional cell with a fixed probability (10 %) and amplitude (~6.5 mV). **(b)** Fine-scale correlations were absent when model neurons were uncoupled by setting the amplitude of the pulse-coupled depolarizing potential to 0 mV. **(c)** Increasing or decreasing the probability of eliciting a pulse-coupled depolarizing potential in the postjunctional cell increases or decreases, respectively, the correlation between the Correlation Index and the difference in firing rate. **(d)** Increasing or decreasing the amplitude of the depolarizing pulse-coupled depolarizing potential increases or decreases, respectively, the correlation between the Correlation Index and the difference in firing rate.

**Supplementary Figure 3.**
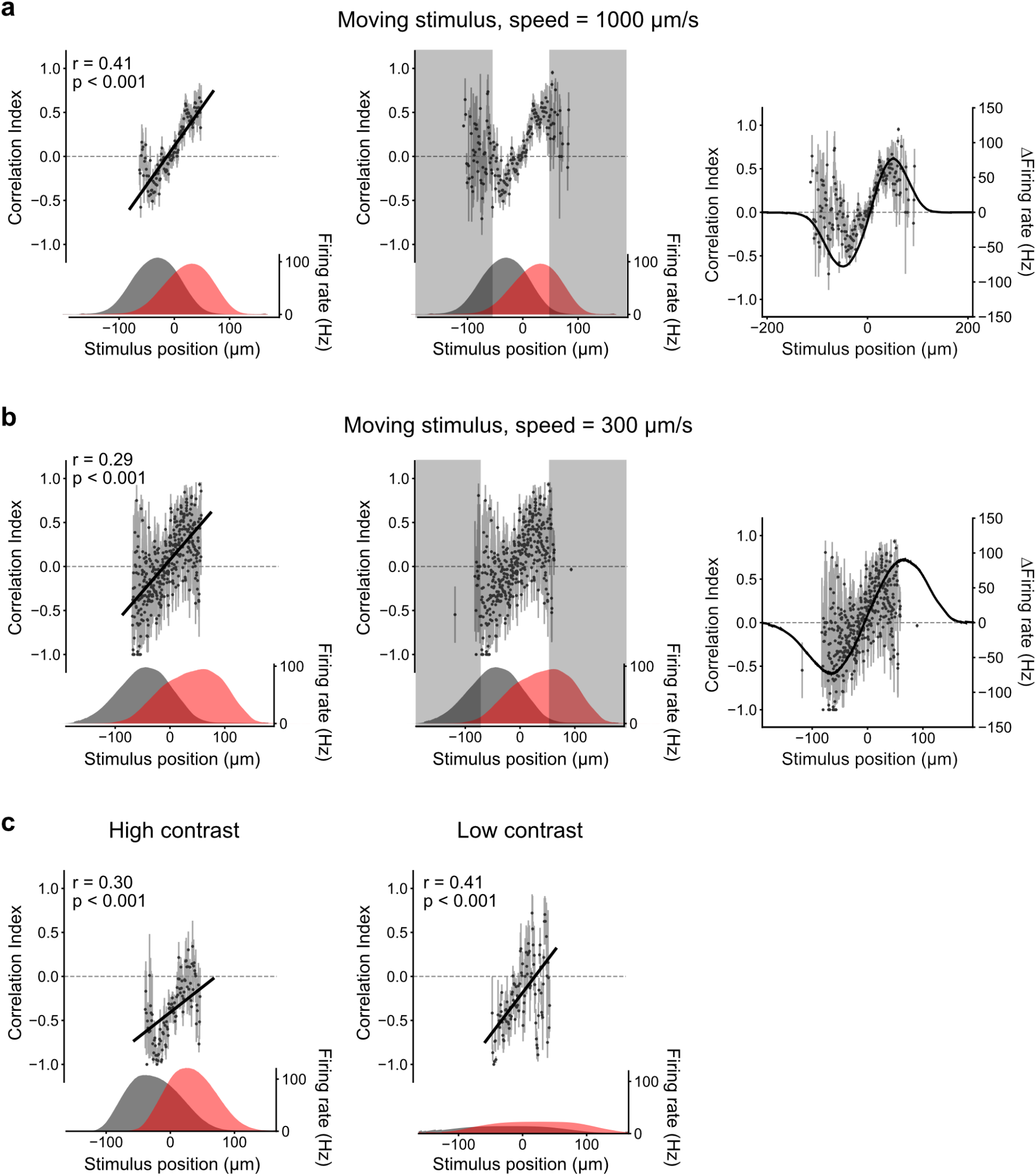
The spike order spatial code is invariant to changes in speed and contrast of the moving stimulus. **(a)** *Left*, Spike order spatial code for fast-moving stimuli (re-plotted from **Figure 2d**, stimulus speed = 1000 μm/s). *Middle*, The Correlation Index was highly variable in regions of low firing rate (grey boxes) due to the low number of spike-pairs. These regions (defined as two standard deviations from the peak firing rate of each cell) were therefore omitted from the analysis included in **Figure 2**. *Right*, CI (black dots) and firing rate difference between the pair of cells (solid black line) are plotted against the stimulus position. The shape of the CI curve mirrors that of the firing rate difference. **(b)** Same as in (a) but for slow-moving stimuli (stimulus speed = 300 μm/s; n = 5 cell pairs). **(c)** The spike order spatial code of fast-moving stimuli (1000 μm/s) at high (*left*) and low (*right*) contrast (contrast was lowered such that the peak spike rate was reduced by roughly 80 %; n = 3 cell pairs; note that the same 3 cell pairs are shown for high and low contrast).

**Supplementary Figure 4.**
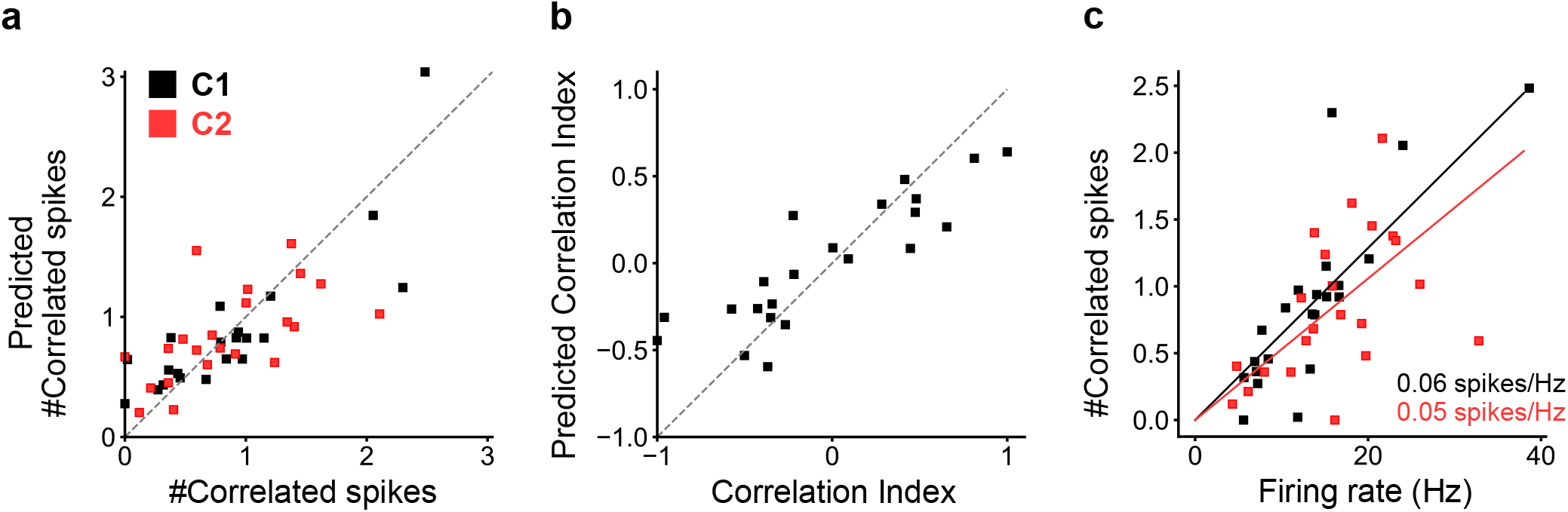
The number of correlated spike-pairs is proportional to the firing rate. Previously it has been shown that spike correlations arise when one coupled ganglion cell fires an action potential, which leads to a backpropagating action potential (bAP), which drives a coupled spikelet in postjunctional dendrites, which then combines with chemical synaptic inputs to drive a dendritic spike, which then drives a postjunctional correlated action potential^2^. As such, increasing the spike rate will increase the number of bAPs, resulting in more coupled spikelets, and this should lead to an increase in the total number of correlated spikes. We tested whether the relationship between spike rate and correlated spikes followed a simple linear relationship, using the following model: *τ* = *p* · *r* where τ is the number of correlated spikes in the postjunctional cell, r the firing rate of the prejunctional cell, and p the proportionality factor ranging from 0 to 1 (note that p can be interpreted as a probability p = τ/r, since 0 ≤ τ ≤ r). We estimated the probability that a spike in one cell will drive a correlated spike in the neighbouring cell (p) as the ratio of the number spikes that caused a correlated spike (τ) and the total number of spikes (r) using the pooled data from all 6 cell pairs (including all stimulus positions) used for **Figure 2a-c**. We then used this probability to predict the number of correlated spike-pairs and the resulting CI based simply on the firing rate of the cells in each pair at each stimulus position. **(a)** The predicted number of correlated spike-pairs is plotted against the observed number of correlated spike-pairs. Predicted and observed values were not significantly different from each other (n = 44, W = 469, P = 0.76, Wilcoxon signed-rank test). **(b)** Predicted CI is plotted against the observed CI. Predicted and observed values were not significantly different from each other (n = 22, W = 115, P = 0.71, Wilcoxon signed-rank test). **(c)** The firing rate of the prejunctional cell was linearly correlated with the number of postjunctional correlated spikes, and this relationship was symmetrical (cell 1, C1: r = 0.49 (Pearson coefficient), P = 0.02; cell 2, C2: r = 0.83 (Pearson coefficient), P < 0.001; P-values were calculated with a permutation analysis of the Pearson correlation).

## Methods

### Whole-mount retinal preparation and electrophysiological recordings

All procedures were performed in accordance with the Canadian Council on Animal Care and approved by the institute’s Animal Care Committee. Retinae of Hb9::eGFP transgenic mice^7^ were prepared as previously described^2^. Recordings were made as previously described^2^. In brief, a 2-photon laser scanning microscope, set to 920 nm, was used to identify eGFP^+^ cells. To record spikes, cells were recorded from in the loose-patch configuration, using 5-10 MΩ patch pipettes filled with Ringer’s solution. Recordings were made using a MultiClamp 700B amplifier (Molecular Devices) and acquired using custom software written in LabVIEW^2^. Light stimuli were produced as previously described^3^. Unless otherwise noted, all stimuli were shown at maximum contrast.

### Cross-correlograms and Correlation Index

To construct the cross-correlogram, we computed the pairwise relative spike time difference from spike trains recorded from two coupled neurons and plotted them in a histogram with 0.5 ms bins. For all cells, Cell 2 was defined as the reference cell, so negative bins indicate that a spike in Cell 1 preceded a spike in Cell 2, and vice versa. Spike time differences were pooled across stimulus trials. Throughout the paper, we pooled ON and OFF responses as we did not observe any qualitative differences in our results if we separately analyzed correlations within ON or OFF responses (**Supplementary Figure 1**). For static flash experiments in **Figure 2**, in order to be able to pool together data across cells (**Figure 2b,c**), for each cell pair we calculated CI vs. stimulus position, fit the data with a linear regression, and then defined the x-axis intercept as 0 μm on the x-axis.

For moving stimuli (**Figure 2d**), Cell 1 was defined as the cell that responded first to a moving stimulus moving in the preferred direction. Spike time differences (i.e. correlated spikes) were computed in a 5 ms-wide window that was slid across the spike train in 1 ms steps. In order to pool together ON and OFF responses, for each pair of cells we first computed the firing rate over time (which for a moving bar is equivalent to firing rate over distance) using a 200 ms moving window (1 ms steps). Next, individually for both ON and OFF responses, we took the area under the curve when both cells were responding and set the peak (i.e. the inflection point) of this co-active region as 0 μm on the x-axis. We found that the Correlation Index (defined below) could not be accurately determined at stimulus positions that elicited very low firing rates, due to the exceedingly low number of correlated spike-pairs (**Supplementary Figure 3**). We therefore restricted our analysis to the region within two standard deviations from the peak firing rate of each cell.

We devised a Correlation Index (CI) to quantify the asymmetry of the cross-correlogram. CI was defined as

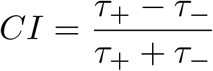

where τ+ and τ− represent the number of spike pairs with spike time difference of 0 to 2 ms and −2 to 0 ms, respectively. To account for stimulus-driven biases, we re-computed spike time difference from randomly shuffled response trials (‘shift predictor’). τ+ and τ− were corrected by subtracting the bias and any negative values were rectified to 0 (**Supplementary Figure 1**). However, the relationship between Correlation Index and difference in firing rate was also present in the ‘raw’ data, before subtracting the ‘shift predictor’ calculated spikes (**Supplementary Figure 1**).

### Model of pulse-coupled neurons

To see if differences in firing rate between pairs of coupled cells were sufficient to account for the modulation of fine-scale correlations, we constructed a simplified computational model that consisted of two adaptive exponential integrate-and-fire neurons (AdEx)^19,20^ with pulsatile excitatory coupling. The voltage (u) evolution is given by

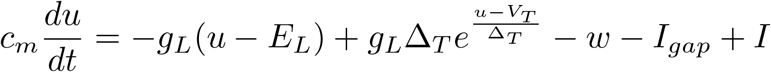

where c_m_ = 1 nF/cm^2^ is the membrane capacitance, g_L_ = 0.3 mS/cm^2^ is the leak conductance, E_L_ = −65 mV is the resting potential, Δ_T_ = 2 mV is the slope factor, V_T_ is the spiking threshold potential, w is a hyperpolarizing adaptation current, I_gap_ is the gap junctional current, and I is the stimulation current. The integration time step was set to dt = 50 μs. V_T_ evolved according to:

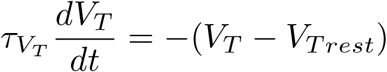

where τ_VT_ = 50 ms is the time constant of the threshold potential and V_Trest_ = −50 mV is the threshold potential at rest. The adaptation current (w) evolved according to:

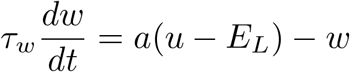

where τ_w_ = 144 ms is the adaptation time constant and a = 4 nS is the adaption current. The gap-junctional current (I_gap_) was modelled as a step current that occurred after an action potential in the other cell. We assigned a fixed probability (transmission probability) such that only a subset of action potentials in one cell drove a correlated spike in the neighbour. Transmission probability was set to 10 % (the effect of increasing/decreasing transmission probability is shown in **Supplementary Figure 2**). The amplitude of the gap junction step current was set so that it would elicit a peak depolarizing potential of ~ 6.5 mV (based on previously measured amplitudes^2^; the effect of increasing/decreasing the amplitude is shown in **Supplementary Figure 2**).

The model neurons were injected with noisy current mimicking synaptic input. The input current was modelled as an Ornstein-Uhlenbeck process^21,22^ with zero mean, variance = 500 pS, and temporal correlation of 4 ms and was adjusted to cause firing rates of 5-40 Hz.

### Data analysis

Throughout the paper, when evaluating the relationship between CI and ΔFiring rate, and CI and stimulus position, we fit the data with a linear regression and present the coefficient of determination r^2^. To further quantify the relationship between CI and ΔFiring rate, and CI and stimulus position, we used the Pearson correlation coefficient (r) and tested its significance using a permutation analysis to evaluate whether the correlation was statistically significant. For the permutation analysis, data points were shuffled, and the Pearson coefficient was calculated, and we repeated this process n-times (n = 10^5^) to generate a null-distribution. The P-value was then defined as the percentile of the observed r value within this distribution (multiplied by 2 for a two-sided test). The performance of the linear model prediction in **Supplementary Figure 4** was evaluated using the Wilcoxon signed-rank test. All data is shown as mean ± SEM.

## Acknowledgements

We thank D. Guitton and A. Krishnaswamy for providing critical feedback on the manuscript. We thank G. Awatramani for discussions on experimental design and feedback on the manuscript. We thank J. Wilsenach for helpful discussions on statistical analyses. We acknowledge funding from the Vision Health Research Network to RT and from the Canada Research Chairs program, the Alfred P. Sloan Foundation, and ONR Global to ST.

## Author Contributions

Experiments were performed by ST. Data was analyzed by RT. Figures were generated by RT. Modeling was performed by RT. The paper was written by RT and ST.

## Data and code availability

Data and custom written codes are available upon request.

## References

1. Connors, B. W. & Long, M. A. Electrical synapses in the mammalian brain. Annu. Rev. Neurosci. 27, 393–418 (2004).

2. Trenholm, S. et al. Nonlinear dendritic integration of electrical and chemical synaptic inputs drives fine-scale correlations. Nat. Neurosci. 17, 1759–1766 (2014).

3. van Welie, I., Roth, A., Ho, S. S. N., Komai, S. & Häusser, M. Conditional Spike Transmission Mediated by Electrical Coupling Ensures Millisecond Precision-Correlated Activity among Interneurons In Vivo. Neuron 90, 810–823 (2016).

4. Trenholm, S. & Krishnaswamy, A. An Annotated Journey through Modern Visual Neuroscience. J. Neurosci. Off. J. Soc. Neurosci. 40, 44–53 (2020).

5. Hidaka, S., Akahori, Y. & Kurosawa, Y. Dendrodendritic electrical synapses between mammalian retinal ganglion cells. J. Neurosci. Off. J. Soc. Neurosci. 24, 10553–10567 (2004).

6. Trenholm, S., McLaughlin, A. J., Schwab, D. J. & Awatramani, G. B. Dynamic tuning of electrical and chemical synaptic transmission in a network of motion coding retinal neurons. J. Neurosci. Off. J. Soc. Neurosci. 33, 14927–14938 (2013).

7. Trenholm, S., Johnson, K., Li, X., Smith, R. G. & Awatramani, G. B. Parallel mechanisms encode direction in the retina. Neuron 71, 683–694 (2011).

8. Trenholm, S., Schwab, D. J., Balasubramanian, V. & Awatramani, G. B. Lag normalization in an electrically coupled neural network. Nat. Neurosci. 16, 154–156 (2013).

9. Thorpe, S. & Gautrais, J. Rank Order Coding. in Computational Neuroscience: Trends in Research, 1998 (ed. Bower, J. M.) 113–118 (Springer US, 1998). doi:10.1007/978-1-4615-4831-7_19.

10. Portelli, G. et al. Rank Order Coding: a Retinal Information Decoding Strategy Revealed by Large-Scale Multielectrode Array Retinal Recordings. eNeuro 3, (2016).

11. Usrey, W. M., Reppas, J. B. & Reid, R. C. Paired-spike interactions and synaptic efficacy of retinal inputs to the thalamus. Nature 395, 384–387 (1998).

12. Carr, C. E. & Konishi, M. A circuit for detection of interaural time differences in the brain stem of the barn owl. J. Neurosci. 10, 3227–3246 (1990).

13. Branco, T., Clark, B. A. & Häusser, M. Dendritic Discrimination of Temporal Input Sequences in Cortical Neurons. Science 329, 1671–1675 (2010).

14. Lien, A. D. & Scanziani, M. Cortical direction selectivity emerges at convergence of thalamic synapses. Nature 558, 80–86 (2018).

15. Trenholm, S. & Awatramani, G. B. Myriad roles for gap junctions in retinal circuits. in Webvision: The Organization of the Retina and Visual System (eds. Kolb, H., Fernandez, E. & Nelson, R.) (University of Utah Health Sciences Center, 1995).

16. Long, M. A., Deans, M. R., Paul, D. L. & Connors, B. W. Rhythmicity without Synchrony in the Electrically Uncoupled Inferior Olive. J. Neurosci. 22, 10898–10905 (2002).

17. Dugué, G. P. et al. Electrical Coupling Mediates Tunable Low-Frequency Oscillations and Resonance in the Cerebellar Golgi Cell Network. Neuron 61, 126–139 (2009).

18. Galarreta, M. Electrical Coupling among Irregular-Spiking GABAergic Interneurons Expressing Cannabinoid Receptors. J. Neurosci. 24, 9770–9778 (2004).

19. Brette, R. & Gerstner, W. Adaptive Exponential Integrate-and-Fire Model as an Effective Description of Neuronal Activity. J. Neurophysiol. 94, 3637–3642 (2005).

20. Clopath, C., Büsing, L., Vasilaki, E. & Gerstner, W. Connectivity reflects coding: a model of voltage-based STDP with homeostasis. 352 (2010) doi:10.1038/nn.2479.

21. Rauch, A., La Camera, G., Lüscher, H.-R., Senn, W. & Fusi, S. Neocortical Pyramidal Cells Respond as Integrate-and-Fire Neurons to In Vivo–Like Input Currents. J. Neurophysiol. 90, 1598–1612 (2003).

22. Köndgen, H. et al. The Dynamical Response Properties of Neocortical Neurons to Temporally Modulated Noisy Inputs In Vitro. Cereb. Cortex 18, 2086–2097 (2008).

